# Extracellular Carbonic Anhydrase Supports Constitutive HCO_3_^−^ Uptake in *Fragilariopsis cylindrus* Regardless of Temperature Changes

**DOI:** 10.1101/2022.09.08.507187

**Authors:** Meng Li, Jodi N. Young

**Author notes:** Corresponding author: Meng Li.

## Abstract

Diatoms, including *Fragilariopsis cylindrus* (*Fcyl*), are the major primary producers in productive polar oceans. Little is known about carbon concentrating mechanisms (CCMs) in polar diatoms and their sensitivity to ocean warming and acidification. Here we characterized the CCM response to temperature in *Fcyl* using Membrane Inlet Mass Spectrometry. *Fcyl* increases RuBisCO expression at lower temperatures to compensate slower catalytic rates but maintains a reliance on HCO_3_^−^ uptake across different temperatures (−2 °C to 9 °C) despite higher CO_2_ solubility at colder temperatures. However, when external carbonic anhydrase (eCA) is inhibited, inorganic carbon usage switches from HCO_3_^−^ uptake to a dependency on CO_2_ diffusion. Incorporating these measurements with modeling, we propose that relying on eCA supported HCO_3_^−^ uptake is an adaptive strategy to the highly dynamic polar ocean environment which experience large fluctuations in [CO_2_] but where HCO_3_^−^ is constantly available.

## Main

Marine photosynthesis accounts for almost half of global biological carbon fixation ^1^. How this carbon sink will respond to rising anthropogenic CO_2_ and warming temperatures has yet to be established. A major confounding variable is the role of carbon concentrating mechanisms (CCMs), biochemical and biophysical cellular mechanisms by which inorganic carbon is elevated around the CO_2_-fixing enzyme, RuBisCO^2^. CCMs enable carbon fixation to be largely insensitive to CO_2_ and instead cells can utilize the much larger pool of HCO_3_^−^ that exists in seawater. CCMs can be energetically costly^3^, and it has been hypothesized that alleviation of the reliance on CCMs could enhance algal growth^4^. However, the complexity of CCMs and their diversity across taxa, coupled with spatial and temporal variation in the marine environment, has raised questions as to whether marine carbon fixation is sensitive to rising CO_2_ and lowering pH^5^.

Polar oceans cover 20% of the global ocean and polar primary production supports ecologically unique ecosystems, drives a strong biological carbon pump and impacts many other large scale biogeochemical processes^6^. Due to extreme environments and remote locations, studying primary production in these environments is difficult. However, these regions are highly sensitive to the effects of rising anthropogenic CO_2_ and already observations have recorded ecosystem regime changes in some polar regions^7,8^.

Primary production in polar oceans appears to be largely insensitive to ocean acidification, attributed to the high concentration of CO_2_ in polar oceans, which results in CO_2_-saturated photosynthesis and a reduced requirement for a CCM^9–11^. However, most polar microalgae possess CCMs, particularly diatoms, which dominate these regions^12–15^. We hypothesize that polar diatoms possess a CCM as an adaptation to the dynamic environment of polar oceans, where strong seasonal cycles, intense blooms and microenvironments within sea ice can result in wildly fluctuating carbonate chemistry, often resulting in low CO_2_ concentrations that could potentially limit fixation by RuBisCO.

Our understanding of CCMs in polar oceans is complicated by the variable effect of temperature on different CCM components. For example, cold temperatures increase CO_2_ solubility, slow diffusion rates, potentially impact membrane permeability, slow chemical (and enzyme) reaction rates and lower half-saturation constants of enzymes. Whether shifting temperatures impact or even unbalance the synergy between different CCM components, including carbonic anhydrases, membrane transporters, pyrenoid structure and RuBisCO kinetics, is yet to be elucidated. In particular, the effect of temperature on extracellular carbonic anhydrase (eCA) could have large impact as the uncatalyzed equilibration between CO_2_ and HCO_3_^−^ is prohibitively slow at cold temperatures. Sea surface temperatures are rising, with the polar ecosystems particularly sensitive^16^. Understanding the temperature impacts on the CCM and carbon fixation in polar oceans will help model and predict the response of polar ecosystems and biogeochemical cycles to the warming ocean.

Here we investigated the CCM response to temperature (−2, 3 and 9 °C) and the impact of eCA in the model polar diatom, *Fragilariopsis cylindrus* (*Fcyl*), a highly productive species in both seawater and sea ice, across both poles. Using Membrane Inlet Mass Spectrometry (MIMS) and an adapted diatom CCM model, we elucidate CCM function in *Fcyl*, particularly the role of eCA in supporting HCO_3_^−^ uptake, across different temperatures.

## Results

### *Fcyl* has eCA activity

eCA activity in *Fcyl* (observed previously by Kranz *et al*.^15^) was confirmed using membrane inlet mass spectrometry (MIMS). O_2_ and isotopic CO_2_ signal traces were used to infer CCM function in *Fcyl*, as described in previous studies for other mesophilic diatom species^15,17,18^. Upon adding *Fcyl* cells into the reaction chamber in the dark (Fig. 1, t1 to t2), the ^18^O exchange between H_2_O and ^18^O-labeled H^13^CO_3_^−^ is accelerated due to carbonic anhydrase activity. Inhibition of eCA with the addition of acetazolamide (AZ) (Fig. 1A, t1 to t2) further accelerates the apparent decline of ^13^C^18^O^18^O and increase of ^13^C^16^O^16^O in the dark, compared with that without AZ (see also Fig. 2A vs Fig. 2B). The effect of AZ on the ^13^C-labeled CO_2_ (and ^12^C-CO_2_) signal traces was also observed during “light on” periods (Fig. 1, t2 to t3 and t4 to t5), demonstrating eCA activity in *Fcyl*.

**Fig. 1.**
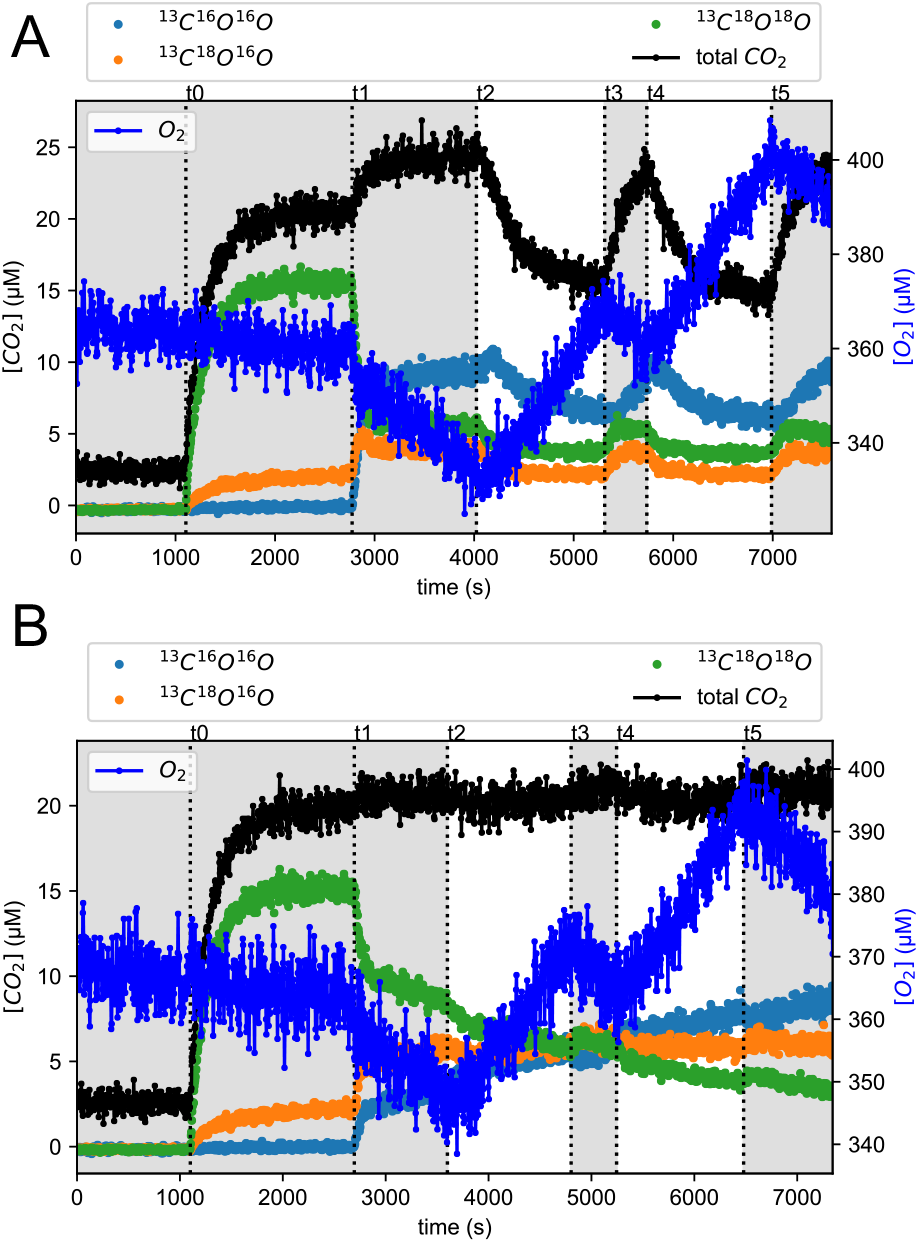
General procedure of *Fcyl* CCM experiments using MIMS. Panel **(A)** and **(B)** present O_2_ (Ar corrected) and CO_2_ signal traces of a typical *Fcyl* CCM experiment at 3°C with and without eCA inhibitor AZ (100μM), respectively. Note that ^12^C^16^O_2_ is not shown but included in total CO_2_ for clarity. The time points t0 through t5 mark the sequential events of adding NaH^13^C^18^O_3_, adding *Fcyl* cells, 1^st^ light on, 1^st^ light off, 2^nd^ light on, and 2^nd^ light off. Shaded areas indicate dark periods in the MIMS reaction chamber.

**Fig. 2.**
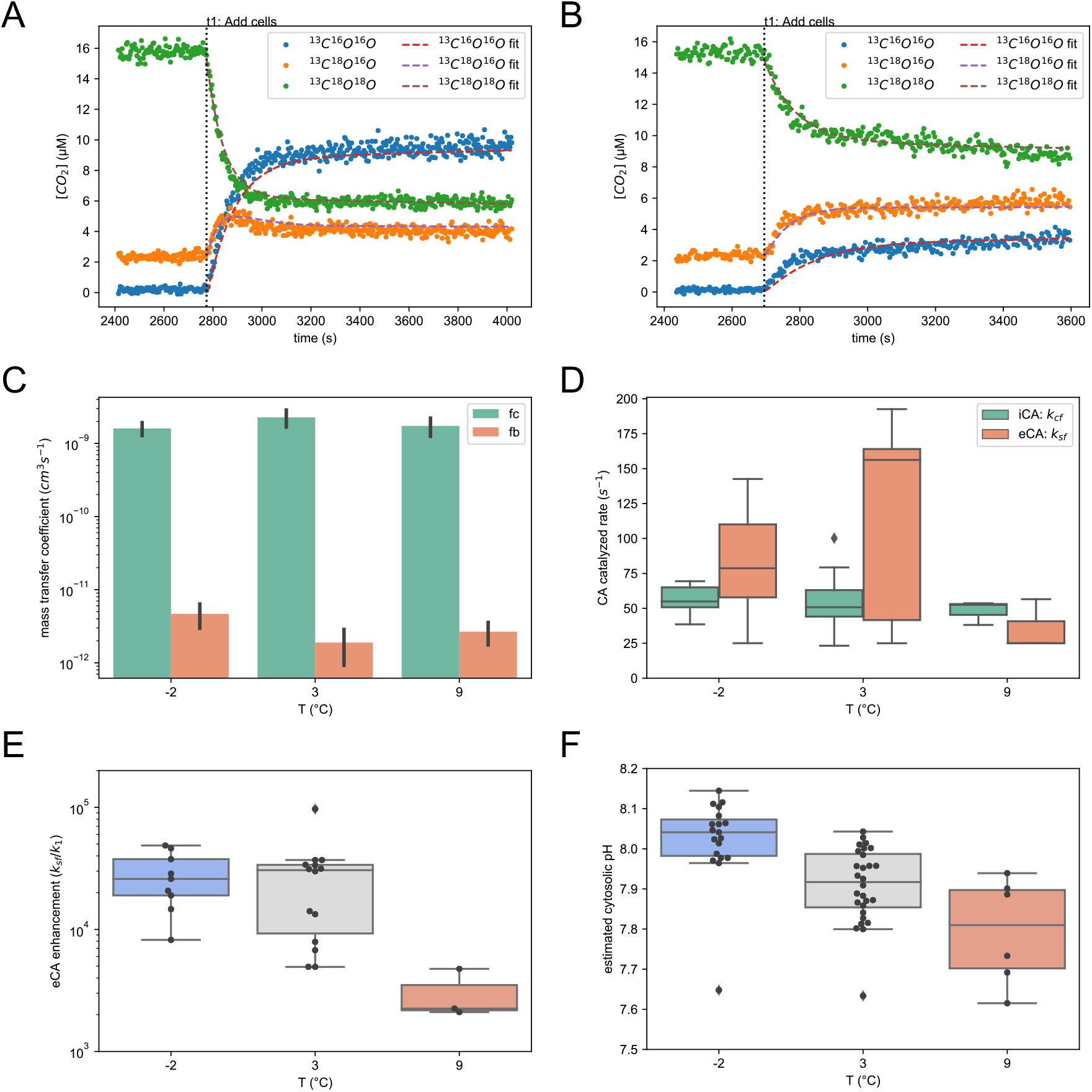
Cell accelerated ^18^O exchange and derived physiochemical parameters. **(A)** and **(B)** examples of ^13^C-CO_2_ signal traces and modeled curve fits after adding cells (t1 to t2 in Fig. 1) with and without eCA inhibitor AZ, respectively. **(C)** Mass transfer coefficients of CO_2_ (f_c_) and HCO_3_^−^ (f_b_) calculated from experiments with the presence of AZ. Error bars represent SD with n = 10, 14, 3 for temperature -2, 3, 9 °C respectively. **(D)** Carbonic anhydrase (CA) activities expressed as catalyzed CO_2_ hydration rates. These values were calculated from experiments without AZ (eCA active) for comparison. One outlier at 3°C for eCA (ksf >400/s) was excluded from the plot for clarity with n = 9, 14, 3 for temperature -2, 3, 9 °C respectively. **(E)** eCA enhancement of CO_2_ hydration rate at cell surface, expressed as *k*_*sf*_ */k*_*1*_. **(F)** Cytosolic pH values estimated from experiments both with and without AZ. Whiskers extend 1.5 times of IQR from box for all box plots.

### *Fcyl* maintains constant eCA activity across different temperatures

^18^O-labeled CO_2_ traces of *Fcyl* cells in the dark (Fig. 2A-B) were used to quantify passive carbon fluxes, i.e., mass transfer coefficients of CO_2_ and HCO_3_^−^ (f_c_ and f_b_), extracellular CA (eCA) activity and intracellular/cytosolic CA (iCA) activity at different temperatures. In agreement with earlier studies on other diatoms^15,17,18^, *Fcyl* cells are more permeable to CO_2_ than HCO_3_^−^, with the magnitude of f_c_, f_b_ estimated around 10^−9^ and 10^−12^ cm^3^ cell^−1^ s^−1^ respectively (Fig. 2C). Calculated membrane CO_2_ permeability (P_c_) at -2 °C was 2.61±0.60 ×10^−3^ cm s^−1^ in this study (Table S1), similar to values reported by Kranz et al. of 2.2×10^−3^ cm s^−1^ at 0°C^15^. Temperature impacts CO_2_ /HCO_3_^−^ diffusion across the cell membrane and boundary layer (Tukey HSD p<0.005) observed for both P_c_ and f_b_ between 3°C and -2°C (Table S1, Fig. 2C).

Absolute values of eCA and iCA activity (s^−1^) were similar across temperatures (Fig. 2D). However, when the catalytic rates of eCA were normalized to uncatalyzed CO_2_ hydration rates, eCA activity enhancement was much higher under lower temperatures (Fig. 2E) due to slower uncatalyzed rates at colder temperatures. Cytosolic pH, estimated from apparent cytosolic equilibrium constant of CO_2_ to HCO_3_^−^ (Eq S11), is temperature dependent, with lower cytosolic pH at higher temperatures, from ∼7.8 at 9 °C to over 8 at -2 °C (Fig 2F). Even though the presence of eCA inhibitor may interfere with the estimated values, the higher cytosolic pH under lower temperature is clear, especially for -2°C (Tukey HSD p≤0.001 compared with 9 °C and 3 °C with AZ, Fig. S2).

### Inhibition of eCA switches carbon uptake from HCO_3_^−^ to CO_2_

*Fcyl* can survive and grow under subzero temperature (−2°C) within sea ice, but faster growth was observed at 3°C without further advantage at higher temperature (9°C) (Fig. S3a). The faster growth rate coincides with faster net O_2_ production rate (Fig. S3a-c). Inhibition of eCA with saturating DIC (∼ 2 mM) did not impact net O_2_ evolution in *Fcyl* (Fig. 3A), which is similar to the low impact of eCA on net photosynthesis observed in other diatoms, e.g. *Thalassiosira* species^18^. However, inhibition of eCA altered inorganic carbon usage, towards CO_2_ versus HCO_3_^−^ uptake from bulk environment (Fig. 3B). With active eCA, *Fcyl* cells primarily used HCO_3_^−^ to support photosynthesis, but switched to CO_2_ usage when eCA was inhibited. This response was insensitive to temperature. Since eCA has been proposed to facilitate CO_2_ supply^18^ or recycle CO_2_^19^ at the surface layer, we also analyzed the CO_2_ /HCO_3_^−^ fluxes through the cytoplasmic membrane during “light on” period (SI Method Eq. S19-24). Compartmentalizing eCA activity to the surface layer and distinguishing surface layer from the bulk environment did not change the observation that *Fcyl* cells use HCO_3_^−^ as the major carbon source for photosynthesis when eCA is active (Figs. 3C and S3d-e). At 3°C, measurements were taken at two salinities (35‰ and 40‰) and there was enhanced CO_2_ leakage from the cells at the higher salinity (t-test p=0.01, Figs. 3C and S3d-e). Net fluxes of CO_2_ hydration/dehydration catalyzed by eCA were calculated (F_CO2eCA_, Eq S25) to demonstrate whether the net role of eCA was to recycle CO_2_ (net hydration) or supply CO_2_ (net dehydration) (Fig. S3f-g). The fraction of CO_2_ supplied or recycled by eCA to total CO_2_ flux across cytoplasmic membrane (F_CO2eCA_ /U_CO2_) was found to be positively correlated with eCA activity (Figs. 3D and S3h).

**Fig. 3.**
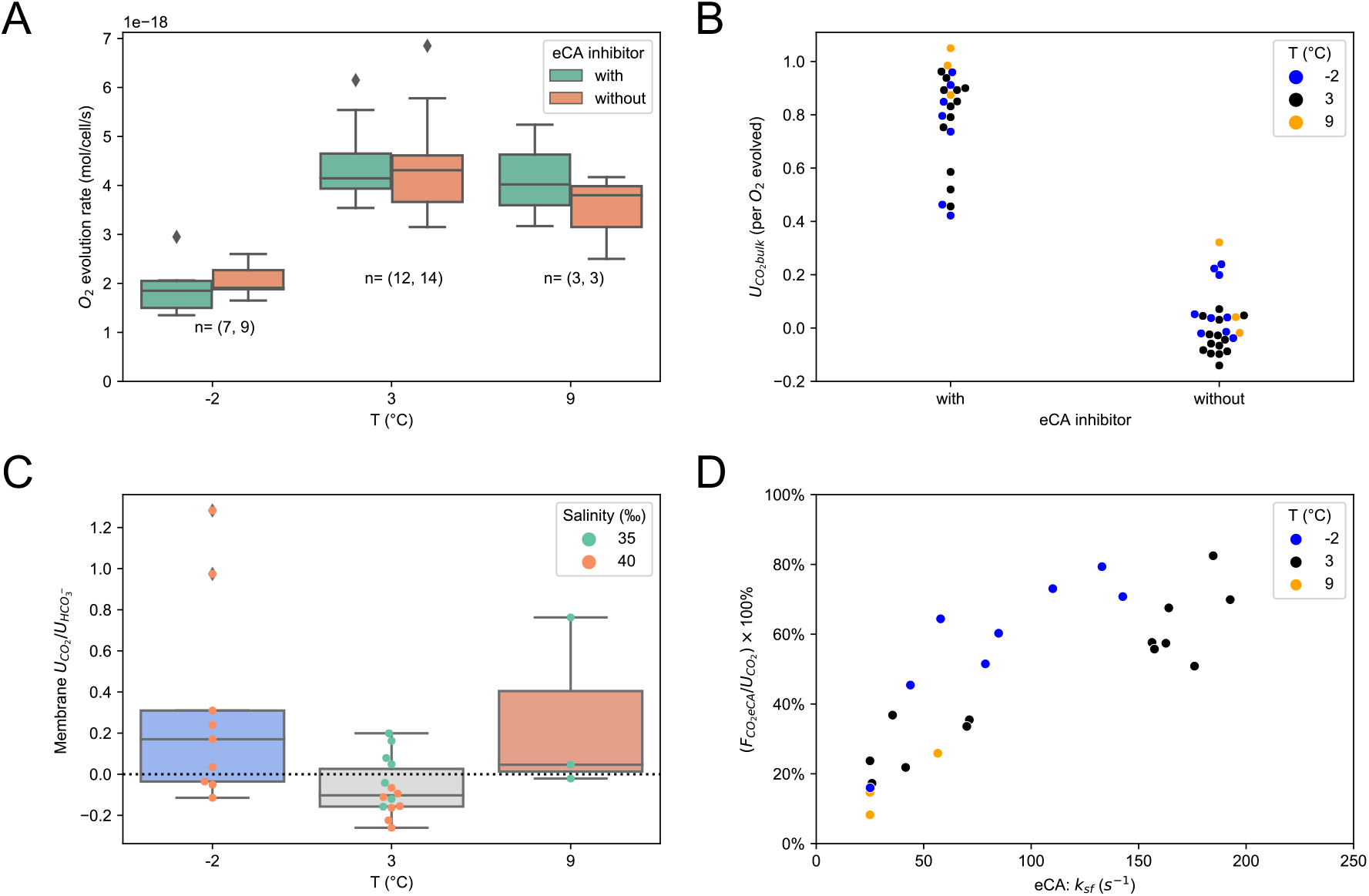
Net photosynthetic rates and DIC usage (CCM) of *Fcyl* in presence/absence of eCA. **(A)** Net O_2_ evolution rate during light on periods at steady state, numbers of biological replicates are denoted under each box plot. **(B)** Steady state bulk CO_2_ uptake (U_CO2bulk_) normalized to O_2_ evolution in the presence or absence of eCA inhibitor AZ, with negative value denoting leaking CO_2_ from cells. **(C)** Without AZ (eCA active), the ratio of CO_2_ uptake (U_CO2_) to HCO_3_^−^ uptake (U_HCO3-_) across plasma membrane. Salinity data of each data point is shown to distinguish the effect of salinity at 3°C. **(D)** The correlation between eCA activity and the percentage of eCA supplied (or recycled) CO_2_ flux to U_CO2_. The percentage of CO_2_ supplied (or recycled) by eCA is defined as the ratio between net CO_2_ flux catalyzed by eCA (F_CO2eCA_) and U_CO2_ (see Fig. S3f-h and Eq S25 for more details).

### Photosynthesis is limited by low DIC/CO_2_ only in the absence of eCA activity

While the presence or absence of eCA does not impact net photosynthetic rate under typical seawater conditions, (i.e., pH 8.10, ∼2 mM DIC), it is required for HCO_3_^−^ uptake. Without eCA activity, *Fcyl* depends on CO_2_ diffusion into cytoplasm to support carbon fixation at RuBisCO. It has been observed that eCA activity is upregulated at low CO_2_ conditions in *Thalassiosira pseudonana*^18^, which suggests eCA activity is important under CO_2_ limitation. To avoid confounding effects of pH, we subjected *Fcyl* cells to lower DIC/CO_2_ (quasi-steady state [CO_2_] <4.5 μM, [DIC] ≤760 μM) conditions at 3°C and analyzed resulting CO_2_ and O_2_ traces for photosynthetic rate and DIC usage (Fig. 4A-B). Under low DIC/CO_2_ conditions, O_2_ evolution was slowed down only when eCA was inhibited (compare t1 to t2 O_2_ traces in Fig. 4A versus Fig. 4B, and see Fig.S4a). Subsequent addition of DIC can restore O_2_ evolution in the presence of eCA inhibitor (t4 to t5 Fig. 4A). This suggests that eCA is required to access the larger pool of DIC (HCO_3_^−^) when CO_2_ is limiting.

**Fig. 4.**
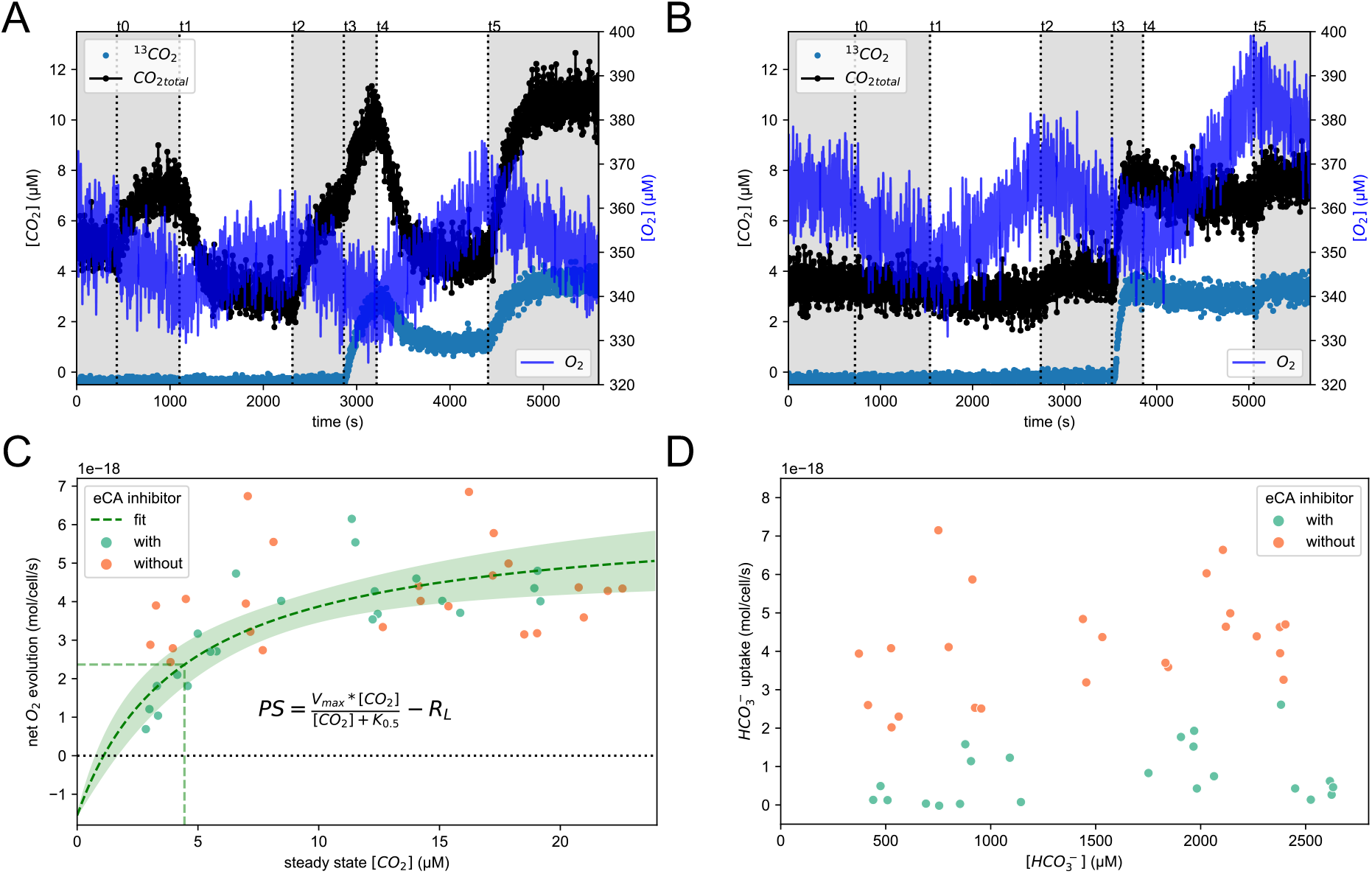
Low DIC/CO_2_ limits *Fcyl* photosynthesis in the presence of eCA inhibitor. **(A)** and **(B)** example O_2_ (Ar corrected) and CO_2_ signal traces of *Fcyl* experiments with low DIC at 3°C in the presence and absence of eCA inhibitor AZ (100μM) respectively. Time points t0 to t5 sequentially mark the events: adding cells, light on, light off, adding ∼400μM NaH^13^CO_3_, light on, light off. Shaded areas indicate dark periods for the MIMS reaction chamber. **(C)** The relationship between photosynthetic rate and steady state extracellular CO_2_ concentration at 3°C. The data combine both saturating DIC and low DIC experiments. The fit curve is based on experimental data with eCA inhibitor. Area within 95% confidence interval of best fit is filled with light green. Vertical line marks the calculated K_0.5_. R_L_ is the light respiration rate, assumed to be equal to the observed mean dark respiration rate (1.55×10^−18^O_2_ mol/cell/s). **(D)** The impact of eCA inhibitor on bulk HCO_3_^−^ uptake rate at 3°C.

The effect of CO_2_ limitation on eCA-inhibited *Fcyl* is clearly demonstrated when net O_2_ evolution rate is plotted as a function of steady state CO_2_ concentration (Fig 4C, with only eCA inhibited cells displaying a Michaelis-Menten response to CO_2_ concentrations). By factoring in respiration (measured during dark steps), the apparent cellular half saturation constant (K_0.5_) of CO_2_ for carbon fixation/gross O_2_ evolution was estimated to be 4.4 ±1.3μM (Fig. 4C), which is close to field measurements of ∼4 μM in polar diatoms communities when eCA was inhibited^15^. This equates to a K_0.5_ for DIC of 750±234 μM in the presence of eCA inhibitor (Fig. S4b). In contrast, *Fcyl* cells with active eCA were not limited by CO_2_ or DIC under our experimental conditions (Figs. 4C and S4b). This indicates that *Fcyl* cells with active eCA can utilize inorganic carbon very efficiently under large range of DIC concentrations and rarely experience DIC supply shortage under natural seawater environment. The concentration of DIC did not impact the preference for inorganic carbon species, with eCA-active *Fcyl* almost exclusively using HCO_3_^−^, while the presence of eCA inhibitor leads to much lower HCO_3_^−^ uptake (Fig. 4D). These results together suggest that eCA activity is required for efficient HCO_3_^−^ uptake across cytoplasmic membrane. For quantitative modeling, we estimated the Michaelis constant K_0.5_ for cytoplasmic membrane HCO_3_^−^ transporter to be ∼400 μM (Fig. S4c).

### RuBisCO kinetics and expression in response to temperature changes

We measured *Fcyl* RuBisCO kinetics in the range of 6∼35°C and fitted the temperature dependence of kinetic parameters (Fig. 5A) of carboxylation turnover rate (*k*_*catC*_) and half saturation constant for CO_2_ in air (K_Cair_). The *k*_*catC*_ of *Fcyl* RuBisCO slows as temperature drops, estimated to be 0.99, 0.55, 0.31 s^−1^ under 9°C, 3°C, and -2°C respectively. With K_Cair_ also dropping with temperature, from 33.5 μM at 9°C to 20.5 μM at 3°C and 13.0 μM at -2°C, there is a slight reduction in *Fcyl* RuBisCO carboxylation efficiency (*k*_*catC*_ /K_Cair_) as temperatures decrease.

**Fig. 5.**
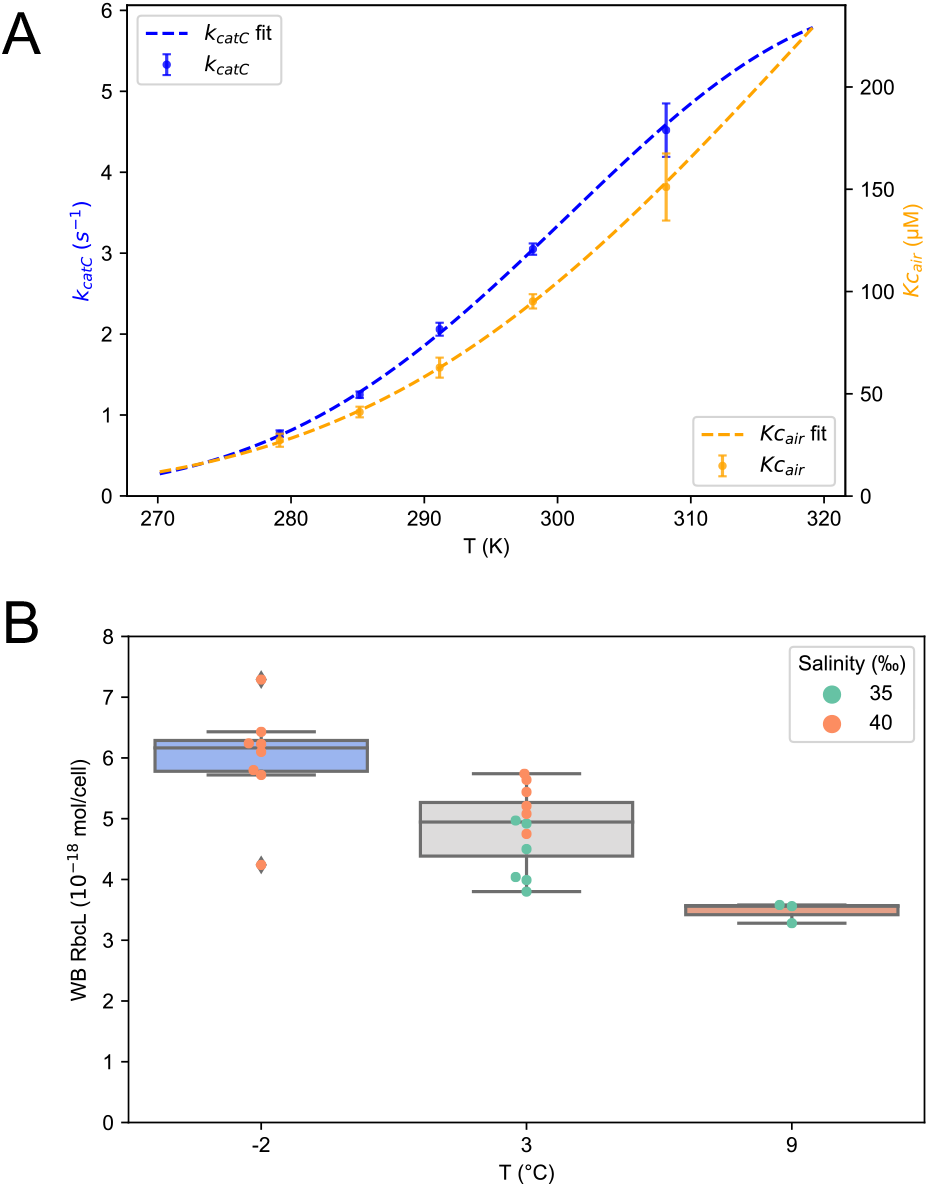
Rubisco kinetic properties and quantities in *Fcyl* at different temperatures. **(A)** Rubisco carboxylation turnover rate (*k*_*catC*_) and CO_2_ concentration at half saturation rate in the presence of air (K_Cair_). The curves were fit according to Eq 2 in Materials and Method. Error bars represent SD of 3 independent measurements. **(B)** Normalized RbcL quantities at different temperatures detected by Western Blot (WB).

The abundance of *Fcyl* RuBisCO protein was upregulated at lower temperature (Figs. 5B, S5). At 3°C, *Fcyl* cells cultured in salinity of 40‰ media had higher RuBisCO content than those in 35‰ (p=0.005) (Fig. 5B). At -2°C, the quantity of RuBisCO per *Fcyl* cell almost doubled (74% more) compared with that at 9°C. These results indicate *Fcyl* cells somewhat compensate lowered catalytic rate of RuBisCO by expressing more RuBisCO in a sea ice (−2°C) environment. Whether all RuBisCO protein at -2°C was fully active is yet to be determined. It is worth noting that Western Blot using Agrisera RbcL standards from spinach or/and our method of extraction underestimated the absolute RuBisCO quantity per cell. That is, the observed O_2_ evolution rates are more than double of the RuBisCO maximum catalytic rates at 3°C. We assume we had incomplete extraction of RuBisCO and 70∼80% saturation^20^, we adjusted the quantified RuBisCO amount by three-fold and used the average quantity at 3 °C for modeling *Fcyl* CCMs at 3 °C.

### Confirming eCA is required for HCO_3_^−^ uptake through modeling

We modeled the carbon fluxes of DIC in *Fcyl* cells by incorporating respiration and updated parameters measured in this study (see Table S2 for details). The model parameters were constrained to describe observed CO_2_ and O_2_ signal traces (Fig. S6). With eCA as the prerequisite for HCO_3_^−^ transport through cytoplasmic membrane, the model shows that under CO_2_ saturating conditions, CO_2_ diffusion and chloroplast HCO_3_^−^ uptake can support sufficient CO_2_ fixation in the pyrenoid when eCA is inhibited (no surface to cytoplasm (SC) HCO_3_^−^ transport) (Fig. 6A). The CO_2_ fixation rates were comparable to those when eCA is active (with SC HCO_3_^−^ transport) (Fig. 6B). To maintain CO_2_ fixation rates with eCA inhibition (no SC HCO_3_^−^ transport), cytosolic CO_2_ concentrations were lower, which drove CO_2_ influx (Fig. 6A), leading to much lower steady state extracellular CO_2_ concentration (10.4 μM) in our closed MIMS system compared with that of eCA intact condition (18.8 μM) (Fig. 6B). This explains why measured total CO_2_ signals decline much faster when eCA inhibitor AZ is present (Figs. 1A-B, 4A-B and S6). To understand the physiological significance of eCA supported HCO_3_^−^ uptake into cytoplasm, we also modeled a diatom bloom situation where pH can reach as high as 8.6^15^. Without eCA (SC HCO_3_^−^ transport), CO_2_ diffusion into the cell requires extremely low cytosolic CO_2_ concentration (∼3 μM) and consequently low cytosolic HCO_3_^−^ concentration (Fig. 6C). This low cytosolic HCO_3_^−^ concentration limits chloroplast HCO_3_^−^ uptake rate and therefore limits the rate of carbon fixation by RuBisCO in pyrenoid. In contrast, *Fcyl* cells modeled with eCA and SC HCO_3_^−^ transport under a diatom bloom condition could sustain high photosynthetic rate (Fig. 6D).

**Fig. 6.**
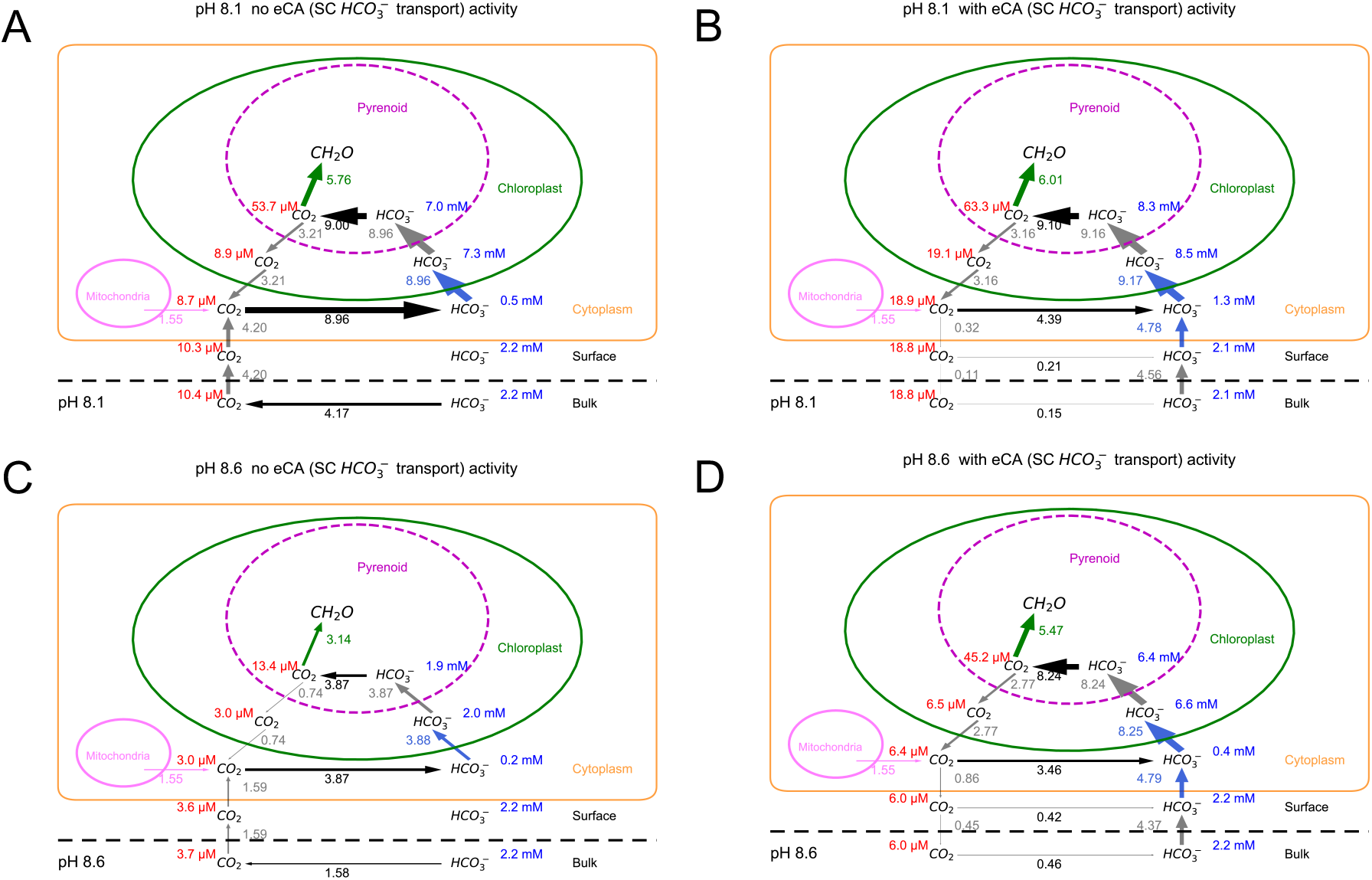
CCM model of inorganic carbon fluxes in *Fcyl* with or without eCA activity at 3°C. eCA is modeled to be the prerequisite for HCO_3_^−^ uptake from surface to cytoplasm (SC HCO_3_^−^ transport). (**A)** and (**B)** is at seawater pH of 8.1, without and with eCA, respectively. (**C)** and (**D)** is at a higher environmental pH (8.6) to elucidate the impact of low CO_2_ during a diatom bloom. Notice the reduced photosynthetic rate and carbon fluxes in panel **(C)** under high pH without eCA activity. Inorganic carbon fluxes (unit in 10^−18^ mol/cell/s) are labeled by the side of arrows, with arrow line width representing relative flux rate. Only net fluxes equal or greater than 10^−20^ mol/cell/s are shown. CO_2_ and HCO_3_^−^ concentrations in different compartments are labeled with concentration unit μM and mM respectively. See SI Method CCM Modeling for details of model description.

## Discussion

The diatom *Fragilariopsis cylindrus* (*Fcyl*) is one of the dominant species in sea-ice phytoplankton communities and polar oceans^21^. The species *Fcyl* has been studied for its temperature acclimation through physiological, genomic, and transcriptomic approaches^22,23^. However, detailed mechanism of how photosynthesis and carbon fixation in *Fcyl* respond to variable environmental conditions is yet to be elucidated. Here, we presented data on inorganic carbon uptake and photosynthesis at varying temperatures including a bottom sea-ice environment, with temperature as low as -2°C.

While eCA activity has been measured in polar diatoms^15^, its role in the CCM and sensitivity to variable temperature had not yet been elucidated. As temperatures lower, carbon uptake into the cell is challenged by slower uncatalyzed rates of CO_2_ hydration/dehydration and reduced diffusive permeability of CO_2_ through surface layer and cytoplasmic membrane. *Fcyl* counters these challenges by regulating both eCA and iCA activity, maintaining high rates of CO_2_ hydration/dehydration that are constant despite changes in temperature. To maintain comparable rates under lower temperatures, *Fcyl* may express more eCA or utilize a cold-adapted eCA. In a global ocean survey, the expression of diatom carbonic anhydrases are generally enhanced under lower temperature^24^.

Under typical seawater conditions (∼2 mM DIC, pH 8.10), the inhibition of eCA does not impact *Fcyl* photosynthetic rates. Similar results of dispensability of eCA under saturating DIC/CO_2_ conditions have been observed in *Skeletonema costatum, Thalassiosira weissflogii, Thalassiosira pseudanana* and *Cylindrotheca fusiformis*^18,25–27^. However, when DIC/CO_2_ is limiting, eCA activity is necessary for higher photosynthetic rates (Figs. 4A-C, S4a). Similar conclusions on the importance of eCA under low CO_2_ or DIC conditions have been also found in other studies of marine phytoplankton^18,25–27^. These studies have proposed that the role of eCA in CCMs is to facilitate the supply of CO_2_ across the cytoplasmic membrane^15,18,28,29^. Trimborn *et al*. also proposed that eCA facilitates the recycling of CO_2_ based on the observations that HCO_3_^−^ uptake makes up over half of the inorganic carbon supply in some diatoms (though these experiment were done with eCA inhibition) under high pH and low CO_2_ conditions^19^. While numerous studies have demonstrated the importance of eCA for the CCM (in some, but not all diatoms), the mechanism by which eCA functions and interpretations had been hindered by the assumption that eCA is only important for the supply (or recycling) of CO_2_.

Using the established MIMS method^17,18,30^ and by quantifying the eCA activity, we have been able to tease apart the inorganic carbon usage of *Fcyl* when eCA is active. Surprisingly, we observed that when eCA is active, *Fcyl* uses primarily HCO_3_^−^ ; in contrast, *Fcyl* mostly depend on CO_2_ diffusion for photosynthesis when eCA is inhibited. The dependence on CO_2_ diffusive supply, i.e., diminished HCO_3_^−^ transport, explains the observed lower photosynthetic rate under low DIC/CO_2_ in *Fcyl* and indicates a HCO_3_^−^ transport system requiring eCA activity. Our hypothesis for an eCA dependent HCO_3_^−^ transport system may also explain similar observations of the necessity of eCA under low DIC/CO_2_ in other diatoms^18,25–27^. This quantitative interpretation of the role of eCA still allows the role of eCA to supply or recycle CO_2_ at the surface layer (periplasmic space)^18,19^, with rate and net direction depending on eCA activity and the cytosolic CO_2_ concentration respectively. Regardless of temperature and salinity, our data indicate that the major role of eCA is facilitating HCO_3_^−^ uptake in *Fcyl* and propose a similar role for other diatoms with eCAs. The function of CA for HCO_3_^−^ transport has been previously demonstrated in the human AE1/CAII system^31^ but has not yet been demonstrated for CCM function.

This study highlights that caution should be taken when interpreting results in the presence of inhibitors alone. In the presence of the eCA inhibitor AZ, our results would agree with previous studies that showed CO_2_ as primary source of entry to the diatom cells^15,25,26^. However, with eCA activity preserved, our results demonstrate HCO_3_ − was the major form of DIC entering *Fcyl* cells. One may raise the concern that eCA activity could mask CO_2_ signals and in turn the analyzed HCO_3_ − usage. To address this question, a simulated CO_2_ signal with active eCA but no cytoplasmic HCO_3_^−^ uptake was compared with our model (Fig. S7). The dashed line in Fig. S7, with no HCO_3_^−^ uptake (or relying on CO_2_ diffusion) but active eCA, cannot explain our observation of little change in CO_2_ signal when eCA is active under illumination. In field research, inorganic carbon usage in marine phytoplankton community is often measured via the disequilibrium assay, where eCA inhibitors are used to eliminate the fast interconversion between ^14^C-labeled HCO_3_^−^ and CO_2_^12,13,15,32,33^. If eCA facilitated HCO_3_^−^ transport plays a significant role within the phytoplankton community, the results obtained from experiments with eCA inhibitors may underestimate the usage of HCO_3_^−13,32^. Methods and data analyses based on the assumption of eCA independent HCO_3_^−^ uptake^15,33^ should be cautioned.

In polar oceans, deep upwelling of CO_2_ and high solubility at cold temperatures can result in CO_2_ concentrations well above that needed to saturate RuBisCO^20^. A CCM or HCO_3_^−^ uptake system seemed unnecessary for phytoplankton to thrive. However, our data demonstrates the usage of HCO_3_^−^ by *Fcyl* regardless of salinities and temperatures tested. This eCA supported HCO_3_^−^ uptake may not be necessary for equilibrated seawater, as supported by the fact that inhibiting eCA generally does not limit photosynthesis under CO_2_ saturating conditions. However, a dynamic polar environment means that CO_2_ is not often at equilibrium in the surface ocean, phytoplankton blooms and sea-ice brine environment can result a pH >9^34,35^ and rapid drawdown of CO_2_ or conversely a buildup on CO_2_ due to winter respiration trapped under sea ice^36^. The eCA supported HCO_3_^−^ transport may be a mechanism for *Fcyl* coping with fluctuating CO_2_ conditions in polar ocean and sea-ice habitats. Changes in pH due to ocean acidification are likely to be small compared to the wide variation experienced seasonally. Likewise, warming of the surface ocean is also unlikely to have a large effect as we observed that *Fcyl* regulates CCM activity to maintain similar function across measured temperatures -2 to 9 °C. In comparison, the Western Antarctic Peninsula has experienced a 1 °C increase in summer surface temperatures since 1950s^37^. However, indirect effects of ocean warming, including loss of sea ice and increased surface stratification, have yet to be elucidated.

## Methods

### Cell culture conditions and growth rate

*Fragilariopsis cylindrus* CCMP1102 (*Fcyl*) cells were cultured in Instant Ocean solution buffered with 25 mM HEPES (pH 8.10) and supplemented with f/2 nutrients^38^ with 0.1 mM silicate. The pH was tuned in accordance with growth temperature and calibrated using thymol blue as described previously^39^. Three different temperatures, -2°C, 3°C, 9°C, were chosen for culture and experiments. Salinity was tuned to 40‰ for -2°C (to prevent freezing and mimic bottom sea ice brine environment) and 3°C, 35‰ for 3°C and 9°C. Both salinities were used at 3°C to disentangle any additional impact of salinity. Light intensity was set at ∼120 μmol m^−2^ s^−1^ at culture bottle surface with illumination from white LED light with 16h: 8h, light: dark cycle. Experimental cells were acclimated to each growth temperature and medium for more than a month before subculturing. Cell densities ranged from 1×10^5^ mL^−1^ at day zero and harvested at cell density between 5×10^5^ mL^−1^ to 1×10^6^ mL^−1^. Growth rates were calculated as

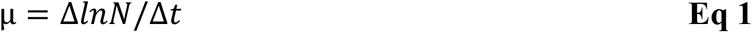

where μ is the growth rate in d^−1^, N is the cell density at given time (day) and t is the time of measurement using Beckman Coulter Z1 cell counter. Time points for growth rates calculations were within three days of the experimental day.

### Membrane Inlet Mass Spectrometry and gas concentration calibrations

Fluxes of isotopically labelled CO_2_ and O_2_ were measured using membrane inlet mass spectrometry (MIMS) using a set up similar to Hopkinson et al. ^17,18^. The cell chamber was cooled to the desired temperature using the water jacket and circulating water bath with ∼15% (v/v) propylene glycol. Each day prior to cell measurements, the MIMS was calibrated similar to Cousins et al.^40^. Calibration for CO_2_ concentration was done by adding known amounts of ^13^C-labeled bicarbonate to 1.2 mL 0.1 N HCl (plus 1M NaCl at -2°C) at a given temperature. ^13^C-CO_2_ mass readings (m/q 45 current) and concentrations (0-327 μM) fit well as a linear relationship (Fig. S1). Factoring the ^13^C/^12^C ratio in the enriched ^13^C-HCO_3_^−^ sample, slopes of linear fits were used for calibrating CO_2_ concentrations later in data processing.

The minimum backgrounds of mass 44 (^12^CO_2_) and 45 (^13^CO_2_) were taken after CO_2_ calibration in HCl by adding 25μL 10 M NaOH. However, we observed that O_2_ concentrations had a slight impact on the signal from these masses, so a further calibration of ^16^O-CO_2_ (mass 44 and mass 45) was done to correct for the impact from O_2_ (mass 32). The O_2_ correction was done by adding Na_2_ S_2_ O_4_ to remove O_2_ in the NaOH solution and assuming a linear relationship (Fig.S1). O_2_ concentrations were calibrated using air saturated water or seawater^41,42^ at a given temperature by sparging for at least 30 min. The minimum background signal of O_2_ was acquired by adding Na_2_ S_2_ O_4_. A linear relationship between O_2_ concentration and signal current was assumed. O_2_ signal was normalized to Ar (mass 40) to correct for air consumption by the MIMS during experimental runs. Conversion of signal current to gas concentrations was calculated using a customized Python script, “MIMS_CO2_calib.py” (available at GitHub: https://github.com/limengwsu/Fcyl_CCM).

### Measuring CO_2_ and O_2_ signals with *Fcyl* Cells

Cells were harvested by centrifugation (5000 g, 2 min) and washed once, followed by pelleting at 10000 g for 1 min, and then resuspended in 25 mM HEPES buffered (pH 8.10) DIC-free artificial seawater (ASW) at growth (same as experimental) temperature. The cells were dark-adapted (>20 min) before added to the MIMS reaction chamber preloaded with buffered ASW with ^18^O, ^13^C-labeled NaHCO_3_ (final concentration ∼2 mM), in the presence or absence of 100 μM acetazolamide (AZ). The final cell density was around 8.6±2.6 × 10^6^ mL^−1^ (mean ±SD) in 1.2 mL and gently mixed via a stir bar (300 rpm). After initial fast gas mixing and the quasi-steady state was reached in the dark (>10 min), a light was turned on (∼250 μmol m^−1^ s^−1^, AmScope 150W halogen bulb) for 20-30 min followed by a light off period. This light on process was repeated with the same cells (Fig. 1). Usually, two experiments, with AZ versus without AZ, were carried out on the same day with samples taken from the same cell culture bottle. For each treatment, at least 3 biological replicates were carried out involving different cells on different days. For low DIC experiments, external ^13^C-NaHCO_3_ was added after first light period (see Results for details).

Fluxes of O_2_ (Ar corrected) in the light and dark measured rates of photosynthesis and respiration. From dark to light transition, diatoms cells need to activate photosynthetic machineries to achieve optimum steady state rate^17^. To avoid confounding impacts of dark adaptation, two “light on” periods (Fig. 1) were measured for steady state photosynthetic rate and CO_2_ signals. The two “light on” periods do not differ in their net O_2_ evolutionary rates (Fig. S3b). Similarly, salinity does little to net O_2_ evolutionary rates (Fig. S3c). In the Results, photosynthetic rates and CCM (carbon usage) were derived from second “light on” periods and 3°C included data from both salinity at 35‰ and 40‰ if not specified.

### Western blot and RuBisCO quantification

*Fcyl* cells were harvested on the day of MIMS experiments and kept in -70 °C before RuBisCO quantification. Cell pellets were resuspended in Tris-Tricine SDS-PAGE sample solubilization buffer^43^ with final cell density of 1.00 × 10^4^ μL^−1^ and 10 μL were loaded for SDS-PAGE followed by Western blot. Western blot transferring was done using BioRad system on PVDF membranes followed by blocking with 3% BSA in TBST. The detection of the large subunit of RuBisCO (RbcL) was done as described in Young et al. ^20^ using global RbcL antibody (Agrisera).

### RuBisCO kinetics

Measurements of *in vitro* RuBisCO kinetics were carried out according to Young et al.^44^. Briefly, RuBisCO active sites were quantified using [^14^C] 2-CABP binding assay as described by Sharwood et al. ^45^. For the ribulose-P_2_ dependent, ^14^CO_2_-fixation assays, crude extracts of soluble diatom protein were incubated with 15 mM NaH^14^CO_3_ and 15mM MgCl_2_ at relevant temperature (6, 12, 18, 25, 35 °C) for 10-15 min to activate RuBisCO. This extract was added to 7 mL septum-capped scintillation vials containing reaction buffer pre-incubated to the desired temperature (0.5 mL of 100mM EPPS-NaOH, pH 8, 15 mM MgCL_2_, 0.6 mM ribulose-P_2_, 0.1 mg ml^−1^ carbonic anhydrase) equilibrated with 21 % (v/v) O_2_ in N_2_ and five differing concentrations of ^14^CO_2_ (between 10∼115 μM; specific activities of ∼1800 cpm per nmol CO_2_ fixed). Values for the half saturation constant for CO_2_ in air (K_Cair_) and maximal carboxylase activity in air (*V*_*Cmax*_) were extrapolated from the data using the Michaelis-Menten equation as described previously^45,46^. The carboxylation turnover rate of RuBisCO (*k*_*catC*_) was calculated by dividing *V*_*Cmax*_ by the number of RuBisCO active sites as determined in [^14^C] 2-CABP binding assay. Temperature dependence of *k*_*catC*_ and K_Cair_ was described by fitting following equation as described by Arcus et al. ^47^:

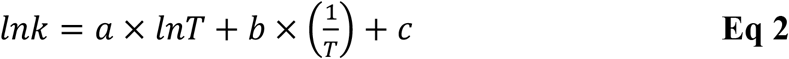

The best-fit constants (a, b, c) were used to extrapolate *k*_*catC*_ and K_Cair_ values at different temperature.

### Data analysis, modeling, and statistical analysis

Gas concentration ([CO_2_] and [O_2_]) over time data were processed using customized Python scripts. The principle of calculating physicochemical parameters, photosynthetic rates and CO_2_ /HCO_3_^−^ have been described by earlier studies^15,17,18,30^. See SI Method Data Analysis for details on calculating *Fcyl* physiochemical parameters, CA activities and carbon usage.

Adapted from “Chloroplast pump” and “Antarctica diatom” models^15,17,48^, we modeled the carbon fluxes of DIC in *Fcyl* cells by incorporating respiration and updated parameters measured in this study (see Table S2 for details). Using CO_2_ drawdown and O_2_ signal traces measured from MIMS, the model used curve fitting estimates of the CO_2_ /HCO_3_^−^ permeability and CA activity within the pyrenoid, and the HCO_3_^−^ uptake activity at chloroplast (Fig. S6).

One major update on those methods was estimating CO_2_ concentration at surface layer and CO_2_ /HCO_3_^−^ fluxes across cytoplasmic membrane when eCA is active (SI Method Data Analysis). Sample data and code for analysis are available on Github (under Fcyl_CCM repository, fit_CO2_Fc.py). The signal processing, modeling, statistical analyses, and curve fitting are largely dependent on the Scipy package in Python^49^ and some data plots were generated using seaborn package^50^. CCM Modeling was done as described in Kranz et al.^15^, with the translation of the MATLAB code to Python script with updated description and parameters (see SI Method CCM Modeling for details).

## Acknowledgements

We would like to thank B. Hopkinson for extensive advice and training for conducting MIMS experiments, and for sharing his mathematical model. We would like to thank S. Whitney for measuring *Fcyl* RuBisCO kinetics in collaboration with JY.

## Author Contributions

JY designed the experiments, ML carried out and optimized experiments, performed data analysis. JY and ML wrote the paper.

## Funding

JY and ML were supported by Simons Foundation Award 561645. JY was also supported by the Sloan Foundation Fellowship and NSF OPP 17445645.

## Code availability

The Python scripts for data analyses and CCM modeling are available on GitHub: https://github.com/limengwsu/Fcyl_CCM

## Notes

### Competing Interest Statement

The authors have declared no competing interest.

