## Supporting Information for "Extracellular Carbonic Anhydrase Supports Constitutive HCO_3_^−^ Uptake in *Fragilariopsis cylindrus* Regardless of Temperature Changes"

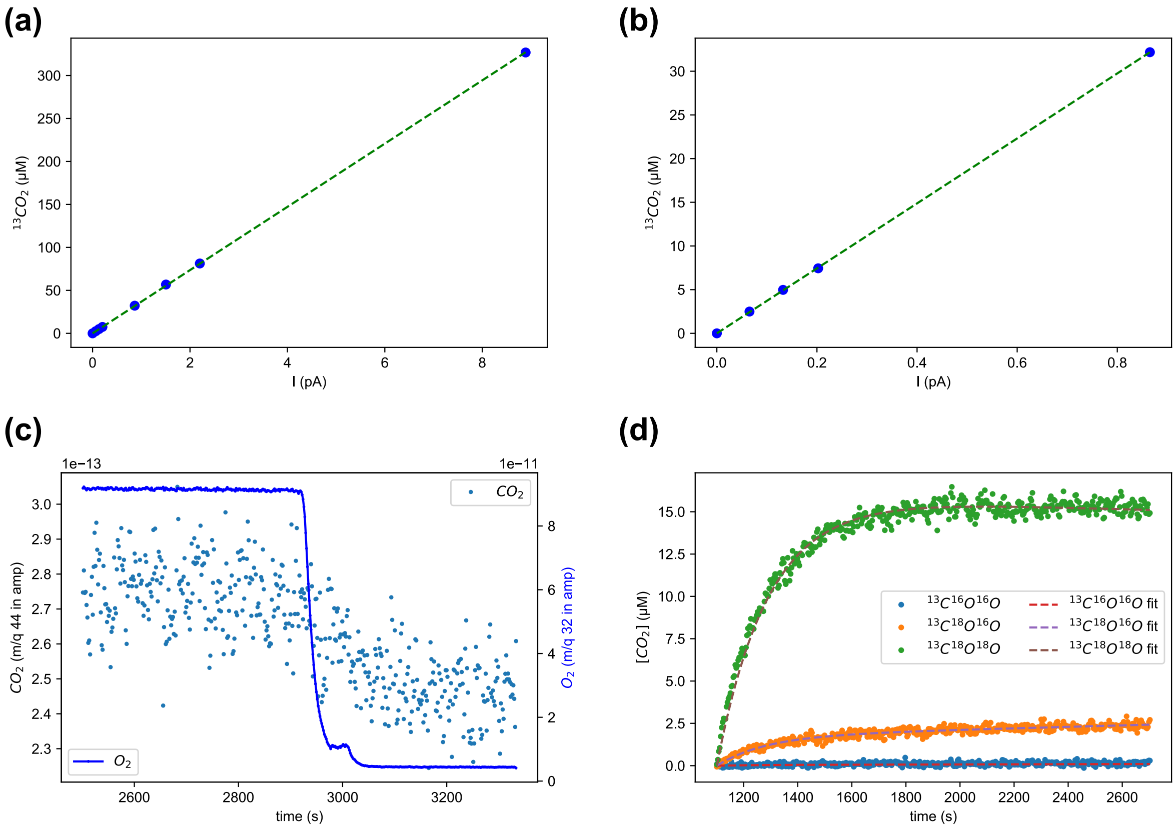


**Fig. S1 MIMS $CO_2$ concentration calibration and uncatalyzed rate constant estimation.** Panel **(a)** and **(b)** present the linear relation between $^{13}CO_2$ signal (m/q 45, I in picoamp) and known $^{13}CO_2$ concentration of a typical experiment at 3°C . Panel **(b)** result is usually used for calculate $CO_2$ concentrations during a CCM experiment. **(c)** The impact of $O_2$ (m/q 32) signal on $^{12}CO_2$ signal. The drop of signals starts from the addition of $Na_2S_2O_4$ to the reaction chamber. The second injection of $Na_2S_2O_4$ solution (around 3000 s) resulted in further decline of $O_2$ signal to real minimum. **(d)** Modeling and curve fitting result of estimating $k_1$ and $k_2$ (see SI Method for details).

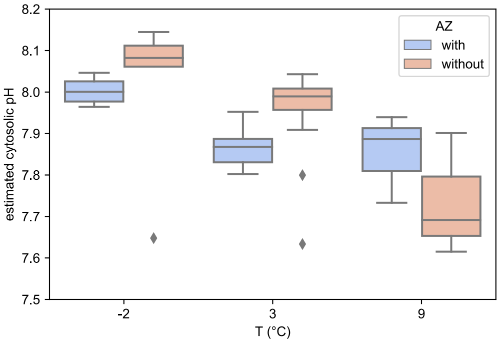


**Fig. S2 The impact of eCA inhibitor (AZ) on estimated cytosolic pH.** The "cytosol" includes everything enclosed by the cytoplasmic membrane, so the cytosolic pH is an overall pH estimated based on "one compartment" model (Hopkinson et al. 2011).

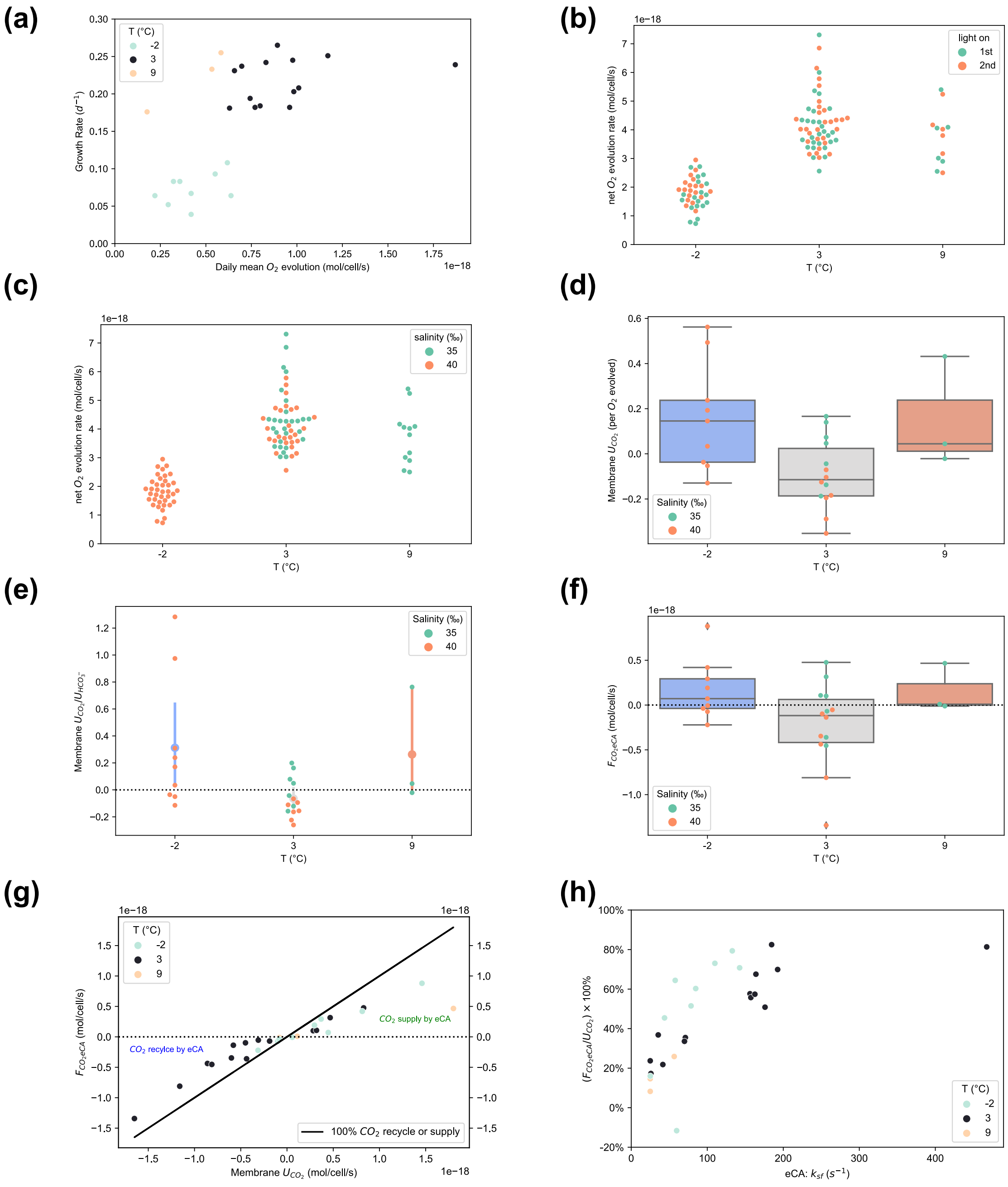

**Fig. S3 Photosynthetic rates and inorganic carbon usage by *Fcyl* during photosynthesis. (a)** Growth rate and mean net photosynthetic ($O_2$ evolution ) rate under different temperature. The daily net $O_2$ evolution rate factors in 2/3 of daylight time and 1/3 of night respiration time. **(b)** and **(c)** present net photosynthetic rate (during the day) in relation to "light on" period and salinity respectively. **(d)** and **(e)** Membrane $CO_2$ uptake ($U_{CO2}$) normalized to net $O_2$ evolution and membrane $HCO_3^-$ uptake respectively. Error bars in **(e)** denote standard deviations. **(f)** and **(g)** Net fluxes from $HCO_3^-$ to $CO_2$ ($F_{CO2eCA}$) at cell surface layer and its relation **(g)** to cytoplasmic membrane $CO_2$ flux ($U_{CO2}$). Negative membrane $U_{CO2}$ indicates leaking of $CO_2$ from cytoplasm, while negative $HCO_3^-$ to $CO_2$ flux indicating recycle of leaked $CO_2$. **(h)** The correlation between eCA activity and the eCA contribution of net DIC fluxes to $CO_2$ supply or recycle (including outliers in **Fig. 3D**).

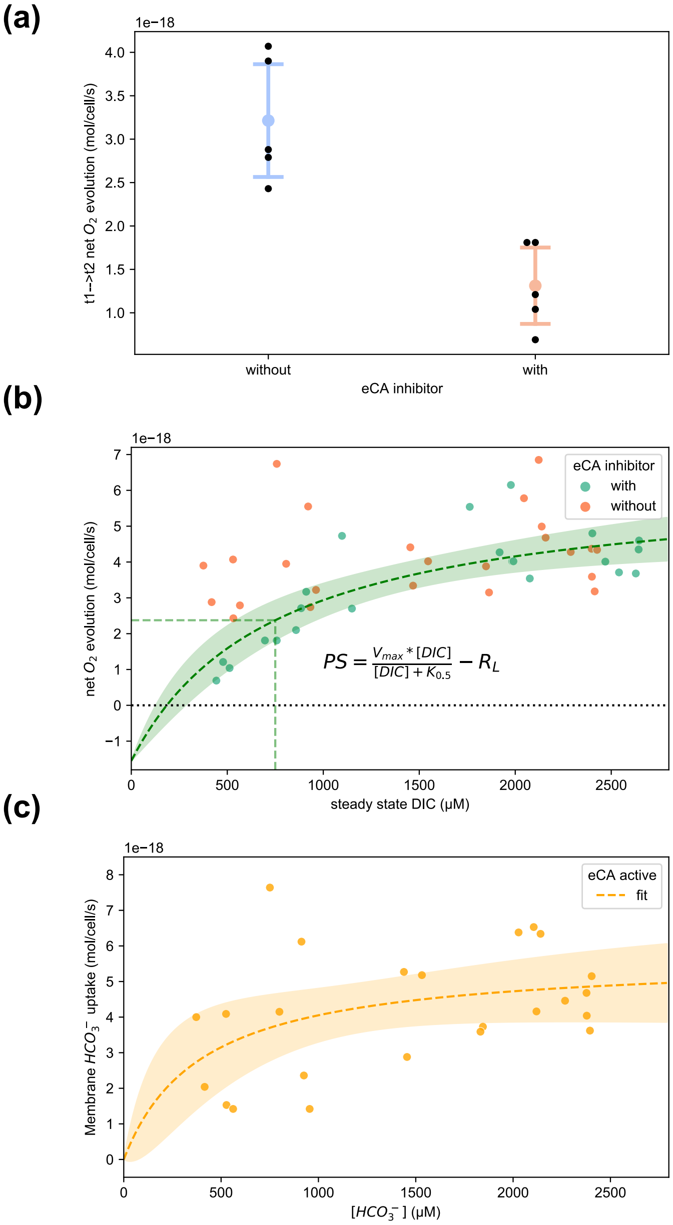


**Fig. S4 The impact of eCA and DIC concentration on photosynthesis and bicarbonate uptake**. **(a)** Net $O_2$ evolution rate with low DIC concentration experiments (Fig. 4A-B t1 to t2) . Error bars present the standard deviations of measured data. **(b)** The eCA activity impacts the dependency of net photosynthetic rate on DIC concentration. Dashed lines show the curve fitting results of the equation shown in the graph for experiments with eCA inhibitor. Areas within 95% confidence interval of best fit is filled light green. Vertical line marks the calculated $K_{0.5}$. $R_L$ is the light respiration rate, assumed to be equal to the observed mean dark respiration rate ($1.55\times10^{-18}$ $O_2$ mol/cell/s). **(c)** The impact of [$HCO_3^-$] on membrane bicarbonate uptake rate at 3°C. The data were fit to Michaels-Menten equation. Area within 95% confidence interval of best fit is filled with light orange.

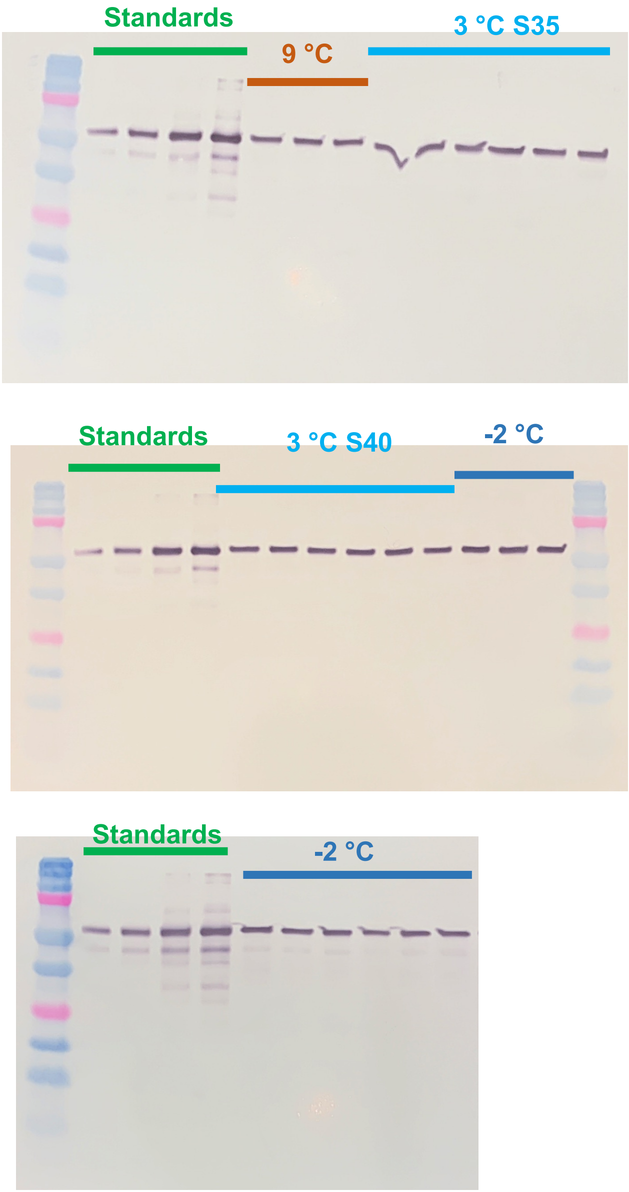


**Fig. S5 Western Blot quantification of RbcL from *Fcyl* whole cell extract**. For each gel, RbcL standards (0.15, 0.30, 0.75, 1.5 pmol) from Agrisera were used for quantification. Sample lanes were marked with *Fcyl* growth conditions. Each lane has loading from 1.00 x $10^5$ cells.

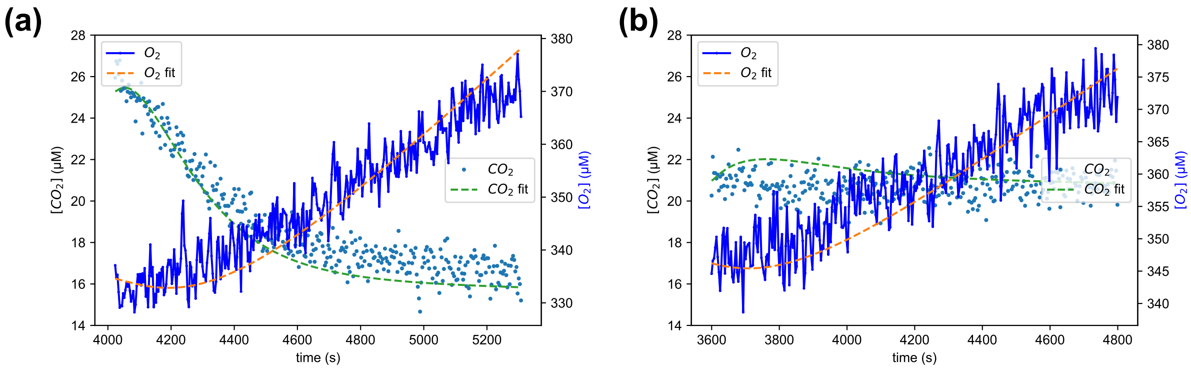


**Fig. S6 Curve fitting results to constrain bicarbonate uptake activity into cytoplasm and chloroplast. (a)** and **(b)** present the modeling and curve fitting of total $CO_2$ and $O_2$ signal for first light period in **Fig. 1A** (eCA inhibited) and **Fig. 1B (**eCA active**)** respectively.

For **(a)**: the best fit values of $V_{max}$ and $K_m$ of chloroplast membrane bicarbonate transporter were $2.92\times10^{-17}$mol/cell/s and 2.22 mM respectively. And the best fit of pyrenoid $CO_2$ hydration rate constant is $6.76\times10^{3}$/s, while the pyr_factor (**Table S2**) is $6.33\times10^{-3}$, i.e., pyrenoid slows $CO_2/HCO_3^-$ diffusion by ~160 fold.

For **(b)**: the best fit values of $V_{max}$ and $K_m$ of chloroplast membrane bicarbonate transporter were $1.97\times10^{-17}$mol/cell/s, and 2.94 mM respectively, indicating different apparent operating kinetics and/or energetic cost when compared with that of **(a)**. The best fit of pyrenoid $CO_2$ hydration rate constant is $5.79\times10^{3}$/s, while the pyr_factor is $6.06\times10^{-3}$, close to the values estimated when eCA is inhibited **(a)**. For modeling purpose, to describe the average photosynthetic rate, these numbers were adjusted as shown in **Table S2**.

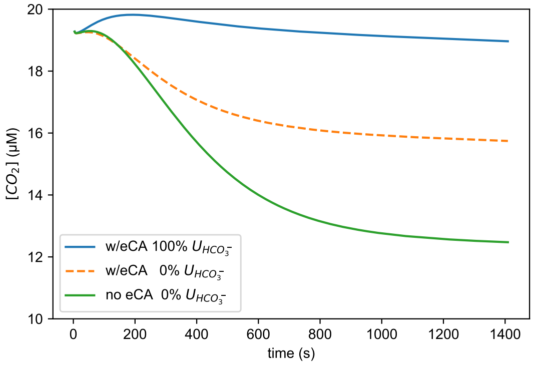


**Fig. S7 The impact of eCA activity on MIMS $CO_2$ signal.** Simulated results of extracellular (bulk) $CO_2$ concentrations that can be detected by MIMS at 3 °C with *Fcyl* cells during "light on" period. To predict the impact of eCA activity without $HCO_3^-$ uptake, the cytoplasmic bicarbonate uptake Vmax, "Vm_Bc" in Table S2, was set to 0 while keeping intact eCA activity in "w/eCA 0% $U_{HCO_3^-}$ ". Note that the simulated dashed line has a significant $CO_2$ signal drop during a typical experimental time span, which should be distinguishable from the solid line of "w/eCA 100% $U_{HCO_3^-}$ " and detected by MIMS. The solid lines present $CO_2$ signal changes similar to our experimental observations in relation to eCA activity.

**SI Method**

**Data Analysis**

Data with gas concentrations were used to calculate physiochemical parameters, photosynthetic rate ($O_2$ evolution), respiration rate ($O_2$ consumption), and CCM carbon fluxes. DIC usage and passive diffusion calculations were based on models described in previous studies (Badger *et al.*, 1994; Hopkinson *et al.*, 2011, 2013; Hopkinson, 2014). Some details and updates are presented below according to Fig. 1.

Kinetics of uncatalyzed interconversion between $CO_2$ and $HCO_3^-$.

The initial $^{13}C$-$CO_2$ (mass 49, mass 47, and mass 45) signals before adding labeled $NaHCO_3$ were set to zero as baseline. By adding $^{13}C$, $^{18}O$ – labeled $NaHCO_3$ (~2 mM final) into DIC-free buffer at certain temperature (t0 to t1 in Fig. 1), the protonation of $HCO_3^-$ leads to the rise of $^{13}C$-$CO_2$ signals. The $^{18}O$-exchange between $H_2O$ and $HCO_3^-$ results in slow decline of $^{18}O$ -labeled $^{13}C$ -$CO_2$. The signals can be used to estimate $k_1$, $k_2$ (or $k_{uf}$, $k_{ur}$ in Hopkinson 2013 et al.), the first order kinetic rate constants of $CO_2$ and $HCO_3^-$ interconversion, by using following differential equations and curve fitting.

$$\frac{d\boldsymbol{C_e}}{dt} = -k_1 \times \boldsymbol{C_e} + k_2 \times \boldsymbol{HB_e}$$

**(Eq S1)**

$$\frac{d\boldsymbol{B_e}}{dt} = k_1 \times \boldsymbol{GC_e} - k_2 \times \boldsymbol{B_e}$$

**(Eq S2)**

Where $\mathbf{C_e}$ and $\mathbf{B_e}$ are $^{13}C$-$CO_2$ and $^{13}C$-$HCO_3^-$ matrices; **G** and **H** are corresponding transformation matrices as described in Hopkinson et al., 2011.

With $^{13}C$-$CO_2$ signal traces, the rate constants, $k_1$ and $K_{eq}$ (apparent equilibrium constant of $CO_2$ hydration, so $\boldsymbol{k_2 = k_1/K_{eq}}$) can be estimated. In the meantime, total added $^{13}C$-labeled [$HCO_3^-$] and fraction of $^{18}O$ in $^{13}C$, $^{18}O$ -labeled $HCO_3^-$ can be calculated (see Github repository "Fcyl_CCM" for details).

Estimating eCA and iCA activities.

Due to the presence of carbonic anhydrases, eCA and iCA, in Fcyl cells, the exchange of $^{18}O$ between $H_2O$ and $HCO_3^-$ can be accelerated. Adding Fcyl cells into MIMS reaction chamber leads to fast drop of $^{13}C^{18}O^{18}O$ (t1 to t2 in Fig. 1). **In the presence of eCA inhibitor AZ (t1 to t2 in Fig. 1A)**, the mass transfer coefficients of $CO_2$ and $HCO_3^-$ ($f_c$ and $f_b$) were estimated using the "One-Compartment Model" (Hopkinson *et al.*, 2011). In the meantime, iCA catalytic activity ($k_{cf}$) and apparent cytosolic $CO_2$ hydration equilibrium ($K_{cyto}$, so $k_{cr} = k_{cf}/K_{cyto}$) were estimated using modeling and curve fitting technique.

With the constraints of $f_c$ and $f_b$, the cytoplasmic membrane mass transfer coefficients for $CO_2$ and $HCO_3^-$, $f_{cM}$ and $f_{bM}$ respectively, were calculated using equations described in Hopkinson *et al.*, 2011.

$$\frac{1}{f_c} = \frac{1}{f_{cM}} + \frac{1}{f_{cBL}}$$

**(Eq S3)**

$$\frac{1}{f_b} = \frac{1}{f_{bM}} + \frac{1}{f_{bBL}}$$

**(Eq S4)**

Where $f_{cBL}$ and $f_{bBL}$ are boundary layer mass transfer coefficients estimated using diffusivity calculated according to Zeebe (Zeebe, 2011). With some updates to the eCA model presented by Hopkinson *et al.*, 2013, the following equations describe the $CO_2$ and $HCO_3^-$ fluxes in the dark after adding cells **without eCA inhibitor** (t0 to t1 in Fig. 1B).

$$\frac{d\boldsymbol{C_e}}{dt} = -k_1 \times \boldsymbol{C_e} + k_2 \times \boldsymbol{H}\,\boldsymbol{B_e} + f_{cBL} \times N \times \frac{\boldsymbol{C_s} - \boldsymbol{C_e}}{V_e}$$

**(Eq S5)**

$$\frac{d\boldsymbol{B_e}}{dt} = k_1 \times \boldsymbol{G}\,\boldsymbol{C_e} - k_2 \times \boldsymbol{B_e} + f_{bBL} \times N \times \frac{\boldsymbol{B_s} - \boldsymbol{B_e}}{\boldsymbol{V_e}}$$

**(Eq S6)**

$$\frac{d\boldsymbol{C_s}}{dt} = -k_{sf} \times \boldsymbol{C_s} + k_{sr} \times \boldsymbol{H}\,\boldsymbol{B_s} + \left(\frac{1}{V_s}\right) \times (f_{c_{BL}} \times (\boldsymbol{C_e} - \boldsymbol{C_s}) + f_{c_M} \times (\boldsymbol{C_i} - \boldsymbol{C_s}))$$

**(Eq S7)**

$$\frac{d\boldsymbol{B_s}}{dt} = k_{sf} \times \boldsymbol{G}\,\boldsymbol{C_s} - k_{sr} \times \boldsymbol{B_s} + \left(\frac{1}{V_s}\right) \times (f_{b_{BL}} \times (\boldsymbol{B_e} - \boldsymbol{B_s}) + f_{b_M} \times (\boldsymbol{B_i} - \boldsymbol{B_s}))$$

**(Eq S8)**

$$\frac{d\boldsymbol{C_i}}{dt} = -k_{cf} \times C_i + k_{cr} \times \boldsymbol{H}\,\boldsymbol{B_i} + \frac{1}{V_c} \times f_{cM} \times (\boldsymbol{C_s} - \boldsymbol{C_i})$$

**(Eq S9)**

$$\frac{d\boldsymbol{B_i}}{dt} = k_{cf} \times \boldsymbol{G}\,\boldsymbol{C_i} - k_{cr} \times \boldsymbol{B_i} + \frac{1}{V_c} \times f_{bM} \times (\boldsymbol{B_s} - \boldsymbol{B_i})$$

**(Eq S10)**

The subscripts for **B** and **C** denote the location of cellular compartment from bulk/extracellular (**e**), surface layer (**s**), and intracellular (**i**). $V_e$ denotes total extracellular volume (~1.2 mL) while $V_s$ and $V_c$ denote individual cell surface volume and cellular volume. $V_c$ was measured using Beckman Coulter cell counter and $V_s$ was estimated by assuming a 0.1 µm surface layer outside cytoplasm. N is the total number of cells added to MIMS reaction chamber. The catalyzed kinetic constant of eCA at surface layer expressed as $k_{sf}$, iCA activity expressed as $k_{cf}$, and cytosolic (cytoplasmic overall) apparent equilibrium constant $K_{cyto}$ (and $k_{cr} = k_{cf}/K_{cyto}$) were estimated by fitting measured $\mathbf{C_e}$ data with model equations above (**Eq S5 to S10**) (Fig. 2B). Note that the kinetic constants $k_{sf}$ and $k_{sr}$ are expressed in unit of $s^{-1}$, which do not include a $V_s$ term as described in Hopkinson *et al.*, 2013.

Surface pH or $CO_2/HCO_3^-$ equilibrium constant was assumed the same as bulk solution. The ratio between $K_{eq}$ and $K_{cyto}$ was used to estimate the apparent cytosolic pH ($pH_{cyto}$).

$$pH_{cyto} = pH_e + \log_{10}\frac{K_{cyto}}{K_{eq}} = 8.10 + \log_{10}\frac{K_{cyto}}{K_{eq}}$$

**(Eq S11)**

Photosynthetic rate and DIC usage.

The calculations of net photosynthetic rate (P), apparent $CO_2$ uptake ($U_{CO2bulk}$), and $HCO_3^-$ uptake ($U_{b\_bulk}$) were the same as described in previous studies (Badger *et al.*, 1994; Hopkinson *et al.*, 2011). When normalized to per *Fcyl* cell, the equations are as follows:

$$P = \frac{d[O_2]}{dt} \times \frac{V_e}{N}$$

**(Eq S12)**

$$U_{CO_2bulk} = (k_2[HCO_3^-] - k_1[CO_2] - \frac{d[CO_2]}{dt}) \times \frac{V_e}{N}$$

**(Eq S13)**

$$U_{b_{bulk}} = P - U_{CO_2bulk}$$

**(Eq S14)**

Since those equations describing $CO_2/HCO_3^-$ uptake were based on uncatalyzed $CO_2/HCO_3^-$ interconversion. The $CO_2/HCO_3^-$ uptake rates can only describe the $CO_2/HCO_3^-$ fluxes into the surface layer. In other words, the $CO_2/HCO_3^-$ fluxes across cytoplasmic membrane need further calculation based on surface layer $CO_2$ and $HCO_3^-$ concentrations if eCA is active. In addition, the estimation of dark [$HCO_3^-$] needs to be adjusted when eCA is active. In Badger *et al.*, 1994, $[HCO_3^-]_{dark}$ was estimated as:

$$[HCO_3^-]_{dark} = \frac{\left\{\left(\frac{d[CO_2]}{dt}\right)_{dark} + k_1[CO_2]_{dark} - respiration\right\}}{k_2}$$

**(Eq S15)**

Which is based on the [$CO_2$] changing rate:

$$\frac{d[CO_2]_{dark}}{dt} = k_2[HCO_3^-]_{dark} - k_1[CO_2]_{dark} - \frac{d[O_2]_{dark}}{dt}$$

**(Eq S16)**

**Eq S15** and **S16** assume that for every $O_2$ consumed, there is one $CO_2$ released into the bulk solution. This is a close estimation when eCA is inactive. However, if eCA is active, it is reasonable to speculate that at least some, if not all, $CO_2$ released will be converted rapidly into $HCO_3^-$. By assuming $CO_2$ released reaches equilibrium with $HCO_3^-$, that is:

$$\frac{d[CO_2]_{respiration}}{dt} = \frac{-d[O_2]_{dark}}{dt} \times \frac{1}{1 + K_{eq}}$$

**(Eq S17)**

So, when eCA is active, the [$HCO_3^-$] was estimated using following modified equation:

$$[HCO_3^-]_{dark} = \frac{\left\{\left(\frac{d[CO_2]}{dt}\right)_{dark} + k_1[CO_2]_{dark} + \frac{d[O_2]_{dark}}{dt} \times \frac{1}{1 + K_{eq}}\right\}}{k_2}$$

**(Eq S18)**

If the term $1/(1+K_{eq})$ is not introduced in **Eq S18**, as in **Eq S15**, the $[HCO_3^-]_{dark}$ will be underestimated because $d[O_2]_{dark}/dt$ is a negative term. The underestimation of [$HCO_3^-$] will lead to underestimation of $U_{CO_2bulk}$, which can generate the misleading conclusion of significant $CO_2$ emission/leaking from *Fcyl* cells when eCA is active. By taking the equilibrium at surface layer into account, the bulk $CO_2$ usage did not surpass $HCO_3^-$ uptake when eCA is active (Fig. 3B).

To "peel" the boundary layer and calculate the $CO_2/HCO_3^-$ fluxes across the cytoplasm membrane, the surface layer $[CO_2]$ and $[HCO_3^-]$ were estimated using boundary layer mass transfer coefficients of $CO_2$ and $HCO_3^-$ ($f_{cBL}$ and $f_{bBL}$).

$$U_{CO_2bulk} = f_{cBL} \times ([CO_2]_{bulk} - [CO_2]_{srf})$$

**(Eq S19)**

$$[CO_2]_{srf} = [CO_2]_{bulk} - \frac{U_{CO_2bulk}}{f_{cBL}}$$

**(Eq S20)**

$$[HCO_3^-]_{srf} = [HCO_3^-]_{bulk} - \frac{U_{b_{bulk}}}{f_{bBL}}$$

**(Eq S21)**

With the estimated surface concentrations of $[CO_2]_{srf}$ and $[HCO_3^-]_{srf}$ at given time window, the $CO_2/HCO_3^-$ uptake through cytoplasm membrane can be calculated as:

$$\frac{d[CO_2]_{srf}}{dt} = k_{sr}[HCO_3^-]_{srf} - k_{sf}[CO_2]_{srf} + \frac{U_{CO_2bulk} - U_{CO_2}}{V_s}$$

**(Eq S22)**

$$U_{CO_2} = U_{CO_2bulk} + (k_{sr}[HCO_3^-]_{srf} - k_{sf}[CO_2]_{srf} - \frac{d[CO_2]_{srf}}{dt}) \times V_s$$

**(Eq S23)**

$$U_{HCO_3^-} = P - U_{CO_2}$$

**(Eq S24)**

The net flux of $CO_2$ catalyzed by eCA at surface layer ($F_{CO2eCA}$ in Fig. S3f-g) is calculated and compared with $U_{CO2}$ to determine whether eCA supplies or recycles $CO_2$ during carbon acquisition.

$$F_{CO_2eCA} = (k_{sr}[HCO_3^-]_{srf} - k_{sf}[CO_2]_{srf}) \times V_s$$

**(Eq S25)**

If eCA supplies $CO_2$, the net flux of $HCO_3^-$ to $CO_2$ conversion ($F_{CO2eCA}$) by eCA will be positive and provide a portion of $CO_2$ flux into the cytosol during carbon acquisition. If eCA plays a role in recycling $CO_2$, the term $F_{CO2eCA}$ will be negative.

Sample data and analysis packages can be found in Github repository "Fcyl_CCM".

### CCM Modeling

The CCM model for Fcyl or Antarctic diatoms was developed in 2014 (Kranz *et al.*, 2015). The original model MATLAB code (under Github repository "Fcyl_CCM/Kranz_2014") was translated into Python script ("Antdiatom_model.py") and adjusted for current study (under Github repository "Fcyl_CCM/Li_2022/Fcyl_CCM.py"). The key updates in current model include (1) adding activation curve for photosynthesis and $HCO_3^-$ transport, (2) factoring in respiration for $CO_2$ fluxes, (3) updates on units and some parameter estimations.

First, the activation of photosynthesis and $HCO_3^-$ transport was described as:

$$f_{active} = 1 - e^{0.045(t-t0)}$$

**(Eq S26)**

The fraction of activated Rubisco or $HCO_3^-$ transporters is $f_{active}$, which is used to describe the observed delay of $O_2$ evolution after turning lights on. Time during light on period and the light on time point are denoted as t and t0 respectively, with units in s. The number 0.045 is empirically optimized for 3°C. So, the Rubisco carboxylation rate is calculated as:

$$P_{gross} = m_{Rubisco} \times k_{cat} \times \frac{[CO_2]_{pyr}}{K_C + [CO_2]_{pyr}} \times f_{active}$$

**(Eq S27)**

Where $m_{Rubisco}$ is the total amount of Rubisco in nmol and $[CO_2]_{pyr}$ is the $CO_2$ concentration at pyrenoid. The calculation of $HCO_3^-$ transport follows similar adjustment by $f_{active}$.

Second, respiration by mitochondria was factored in by assuming $CO_2$ released from mitochondria reaches cytosol first, so that:

$$\frac{d[CO_2]_{cyt}}{dt} = -k_{cf} \times [CO_2]_{cyt} + k_{cr} \times [HCO_3^-]_{cyt} + \frac{1}{V_c} \times \{f_{c_M} \times \left([CO_2]_{srf} - [CO_2]_{cyt}\right) + f_{cP} \times \left([CO_2]_{chp} - [CO_2]_{cyt}\right) + R_{mito}\}$$

**(Eq S28)**

Subscripts of $[CO_2]$, "cyt, srf, chp", denote cytosolic, surface layer, chloroplast. $[CO_2]$ is the total concentration in each compartment with unit modeled in µM. $R_{mito}$ and $f_{cP}$ denote the respiration rate (nanomole/s/cell) and the $CO_2$ mass transfer coefficient of chloroplast respectively.

Third, the units in current model equations are in µM, $cm^3$, nanomoles ($10^{-9}$ mol). Also, other updated parameters can be found in Table S2.

Pyrenoid radius is estimated by assuming 628 µM Rubisco octamer in pyrenoid according to Rosenzweig *et al.* (Rosenzweig *et al.*, 2017), so

$$R_{pyr} = \left(\frac{m_{Rubisco}}{8 \times 6.28 \times 10^{-4} mol/L} \times \frac{0.75}{\pi}\right)^{\frac{1}{3}} \times \frac{10^5 \mu m}{dm}$$

**(Eq S29)**

At 3°C the radius $R_{pyr}$ is estimated to be 0.88 µm.

$$f_{cy} = f_{co2_{diffusion}} \times pyr_{factor} = 4 \times \pi \times R_{pyr} \times D_{CO_2} \times pyr_{factor}$$

**(Eq S30)**

where $D_{CO2}$ is the diffusivity of $CO_2$, and $f_{CO2diffusion}$ is the $CO_2$ mass transfer coefficient calculated based on diffusivity of $CO_2$. The $pyr_{factor}$ restricts the diffusion of $CO_2$ and $HCO_3^-$ alike.

**SI Tables**

**Table S1 *Fcyl* mass transfer coefficients and calculated permeability $P_c$ (mean±SD).**

| **Tc** | **-2 °C (n=7)** | **3 °C (n=12)** | **9 °C (n=3)** | **0 °C (Kranz et al. 2015)** |
|---|---|---|---|---|
| $f_c$ ($10^{-9}$ $cm^3$/cell/s) | 1.49±0.29 | 2.30±0.64 | 1.73±0.59 | 24 |
| $f_b$ ($10^{-12}$ $cm^3$/cell/s) | 4.58±2.14 | 1.90±1.06 | 2.66±1.10 | 18 |
| $P_c$ ($10^{-3}$ cm/s) | 2.27±0.45 | 3.86±1.01 | 2.68±0.77 | 2.2 |

Tukey HSD test only found significant difference between -2°C and 3°C, with p-value equals 0.016, 0.004, and 0.002 for $f_c$, $f_b$, and $P_c$ respectively.

### Table S2 *Fcyl* CCM model parameters at 3°C

| Parameters | eCA inhibition | eCA active | Units | Notes | Definition |
|---|---|---|---|---|---|
| N | 1.00E+07 | 1.00E+07 | cells | total cell number | Cell number |
| Ve | 1.00E+00 | 1.00E+00 | $cm^3$ | total bulk volume | extracellular/bulk volume |
| Vc | 4.16E-11 | 4.16E-11 | $cm^3$ | per cell, measured | cytoplasmic volume |
| Vp | 1.04E-11 | 1.04E-11 | $cm^3$ | 25% of Vc | chloroplast stroma volume |
| $r_{bl}$ | 1.00E-01 | 1.00E-01 | µm | | thickness of the surface layer |
| $N_{pyr}$ | 1.00E+00 | 1.00E+00 | | | number of pyrenoids per cell |
| $R_{pyr}$ | calculated | calculated | µm | **Eq S28** | radius of pyrenoid |
| temp | 3 | 3 | °C | | Temperature in Celsius |
| S | 3.50E+01 | 3.50E+01 | ‰ | | Salinity |
| DIC | 2230 | 2230 | µM | | Total dissolved inorganic carbon |
| pHe | 8.10 | 8.10 | | | extracellular pH |
| pHc | 7.87 | 7.95 | | estimated | cytoplasmic pH |
| pHp | 8.15 | 8.15 | | | chloroplast pH |
| $k_{uf}$ | 5.10E-03 | 5.10E-03 | $s^{-1}$ | $k_1$ | uncatatlyzed $CO_2$ hydration rate |
| $k_{ur}$ | 4.41E-05 | 4.41E-05 | $s^{-1}$ | $k_2$ | uncatatlyzed $HCO_3^-$ dehydration rate |
| $k_{sf}$ | 5.10E-03 | 121 | $s^{-1}$ | MATLAB model unit $cm^3/s$ | eCA activity |
| $k_{cf}$ | 1.10E+02 | 1.00E+02 | $s^{-1}$ | | cytoplasmic $CO_2$ hydration rate (CA catalyzed) |
| $k_{pf}$ | 5.10E-03 | 5.10E-03 | $s^{-1}$ | equals $k_{uf}$ | chloroplast stroma $CO_2$ hydration rate |
| $k_{yf}$ | 6.76E+03 | 6.76E+03 | $s^{-1}$ | Hopkinson 2011 $1.8x10^4$ | pyrenoid $CO_2$ hydration rate (CA catalyzed) |
| $k_{cat}$ | extrapolated | extrapolated | $s^{-1}$ | Or $k_{catC}$ | Rubisco carboxylation turnover rate |
| $K_C$ | extrapolated | extrapolated | µM | Or $K_{Cair}$ | Rubisco half saturation constant for $CO_2$ |
| $m_{Rub}$ | 1.45E-17 | 1.45E-17 | mol/cell | $m_{Rubisco}$ | Rubisco content estimated by net $O_2$ evolution rate |
| Vm_Bc | 0.00E+00 | 5.67E-18 | mol/cell/s | eCA impact | maximal $HCO_3^-$ uptake rate into cytoplasm |
| Km_Bc | 4.00E+02 | 4.00E+02 | µM | | half-saturation constant for $HCO_3^-$ uptake into cytoplasm |
| Vm_Bp | 2.92E-17 | 1.97E-17 | mol/cell/s | Higher pH uses "eCA inhibition" numbers | maximal $HCO_3^-$ uptake rate into chloroplast |
| Km_Bp | 1.11E+03 | 1.50E+03 | µM | Higher pH uses "eCA inhibition" numbers | half-saturation constant for $HCO_3^-$ uptake into chloroplast |
| fc | 2.42E-09 | 2.42E-09 | $cm^3\ s^{-1}$ | | |
| fb | 2.11E-12 | 2.11E-12 | $cm^3\ s^{-1}$ | | |
| fc_sm | calculated | calculated | $cm^3\ s^{-1}$ | or denoted as $f_{cM}$ | $CO_2$ mass transfer coefficient from surface to cytoplasm |
| $f_{cp}$ | calculated | calculated | $cm^3\ s^{-1}$ | based on diffusivity of $CO_2$ | $CO_2$ mass transfer coefficient from cytoplasm to chloroplast |
| $f_{cy}$ | calculated | calculated | $cm^3\ s^{-1}$ | **Eq 29** | $CO_2$ mass transfer coefficient from chloroplast to pyrenoid |
| pyr_factor | 6.33E-03 | 6.33E-03 | | **Eq 29** | Ratio between $f_{cy}$ and $f_{CO2}$ by diffusion, 0~1 |
| $R_{mito}$ | 1.55E-18 | 1.55E-18 | mol/cell/s | | Mitochondrial respiration rate under light |
